# Soil legacies of extreme droughts enhance the performance of invading plants

**DOI:** 10.1101/2023.01.30.526304

**Authors:** Madhav P. Thakur, Maartje A. van der Sloot, Rutger A. Wilschut, S. Emilia Hannula, Freddy ten Hooven, Stefan Geisen, Casper W. Quist, Katja Steinauer, Wim H. van der Putten

## Abstract

Extreme droughts can weaken the biotic resistance of native plant communities against the establishment of invading plants. However, we know little about the underlying mechanisms. Using a plant-soil feedback approach, we tested how an extreme drought event alters the soil-mediated biotic resistance of resident native plant communities against invading plant species from native and non-native ranges, namely non-resident natives, native range-expanders, and alien plants. We show that all three types of invading plants performed better in soils with a legacy of extreme drought independent of resident native plant diversity. Path models revealed that extreme drought effects on non-resident natives were mediated by the root biomass of resident native plants and endophytic fungal pathogens during drought, whereas alien plant performance was mediated only *via* the root biomass of resident native plants also during drought. Our results highlight that the performance of resident native plants during extreme drought and subsequent effects on soil fungi determine the performance of invading plants from native and non-native origins after extreme droughts.

## Introduction

The success and failure of invading plants in new environments have intrigued ecologists for a long time^1–4^. Several factors ranging from biotic to abiotic features of the new environment, and from ecological to evolutionary characteristics of the invading plant determine its success^5–9^. One of the classic ideas that have persisted over several decades is how the diversity of native plant communities constrains or facilitates the establishment of invading plants - the so-called diversity-invasibility relationship^2,10–12^. The contemporary understanding of diversity-invasibility relationships is that greater diversity of native plants at local scales (plot level) constrains the establishment of invading plants (e.g., particularly the alien plants that come from other continents), whereas greater diversity of native plants at regional scales (e.g., landscape level observational studies) correlates positively with the diversity of non-native plants^13,14^. Yet, even at the local scale, studies have shown variable biotic resistance of native plant diversity against invading plants, underlain by mechanisms that remain little understood to date^15,16^.

The key idea behind negative diversity-invasibility relationships is that greater plant diversity can utilize resources (e.g., soil nutrients) more efficiently than low-diverse plant communities, which in turn constrains the establishment of invading plants due to the lesser availability of resources^2,17,18^. Recent advances in invasion biology further point to the importance of soil microorganisms like fungi to determine the establishment success (or failure) of invading plants^19–22^. For instance, pathogenic soil fungi in a new environment of invading plants often weakly inhibit them compared to microorganisms from their previous environments^23–25^. Diverse native plant communities could also influence the establishment of invading plants through their ability to steer soil microorganisms^26^. For instance, a recent experimental study showed that greater fungal diversity associated with native diverse plant communities could constrain the performance of alien plants^27^, as such diverse communities are more likely to contain plant pathogens able to infect a new host (i.e., sampling effects). Conversely, pathogenic soil fungi specialized on certain host plants may also become diluted in diverse plant communities^28–30^, which potentially increases the chance of the successful establishment of invading plants given the reduced pathogenic load^31^. Taken together, the performance of invading plants in diverse resident plant communities could very well depend on the type of biotic interactions (e.g., pathogenic, or mutualistic) that emerge between the invading plant and the resident soil microorganisms associated with resident plants.

Diverse native plant communities are further shown to be resistant to external disturbances, such as drought^32–34^, which could affect the success of invading plants^35,36^. With the increasing magnitude and frequency of droughts across the biosphere, it is likely that biotic resistance (or facilitation) of diverse native plant communities through soil microorganisms against invading plants would change. This expectation is plausible as droughts shift the structure and function of soil microbial communities^37–42^, which could subsequently affect the performance of incoming invading plants through drought legacies in the soil^43^.

Here, we study how the performance of three types of invading plants-local native plants, range-expanding (or intracontinental or neonative) plants and alien (or exotic or intercontinental) plants^44,45^ are affected by soil legacies of native plant diversity and extreme drought (Figure 1). Using three broad categories of plant invaders facilitates our understanding of how plant communities may change after extreme drought events, and whether diversity-invasibility relationships vary for various kinds of invader plants. Most diversity-invasibility plant studies have focused on alien plants^46^, whereas local natives (those not yet present in a native community but in near surroundings: hereafter non-resident natives for brevity) and range-expanding plants have rarely been used to examine their potential to invade a new environment that may have the history of native plant diversity and extreme drought.

**Figure 1:**
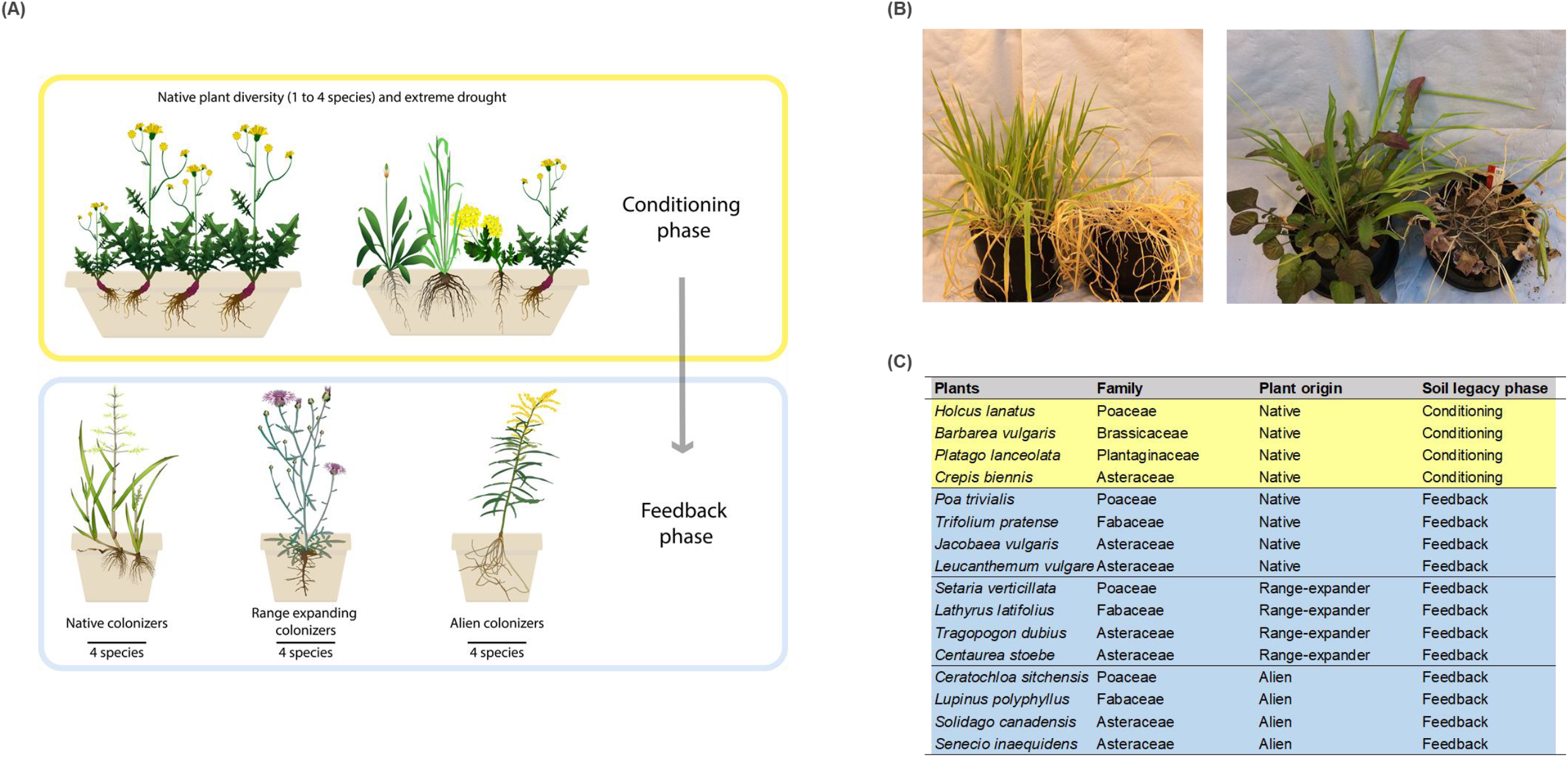
**(A)** Experimental schema showing the conditioning phase with native plant diversity crossed with extreme drought, and feedback phase where those soils were used to grow three groups of invading plants (non-resident natives, range-expanding and aliens) each containing four plant species and grown separately. **(B)** Examples of the extreme drought treatment effects at the end of the conditioning phase: the left image is a monoculture of *Holcus lanatus* with permanent wilting in extreme drought conditions (right side in the image) compared to the one in ambient water conditions, whereas the right image is a mixed plant community with 4 species exposed to ambient and extreme drought treatments (right side in the image). **(C)** List of the plant species used during the conditioning and feedback phase of the experiment.

We use a plant-soil feedback (PSF) experimental approach to investigate soil legacy effects on the performance of invading plants. The PSF approach is a robust way to test the effects of soil microorganisms and soil nutrients in determining the performance of plants through soil legacy or carry-over effects: PSF consists of a conditioning phase (to create a soil legacy) and a feedback phase (to test soil legacy effects, Figure 1)^47,48^. In line with the diversity-invasibility hypothesis, we hypothesize that native plant diversity will constrain the establishment of all three types of invaders, whereas extreme drought will weaken the native plant diversity effect. Moreover, we suspect that these native diversity and drought effects will be mediated *via* soil microorganisms and soil nutrient availability. More specifically, we investigate the role of various functional groups of soil fungi that could affect the performance of invading plants. Given that climate change continues to alter local plant community composition through both local diversity losses and gains, our study attempts to provide insights into underlying mechanisms behind the success and failure of invading plants by zooming in on the roles of soil fungi and soil nutrients.

## Results

The performance of all invading plants was strongly affected by the soil legacy of extreme droughts, whereas the effects of soil legacies of native plant diversity on all three types of invader plants were weak (Table 1, Figures 2 and 3). More specifically, we found that all three types of invading plants had greater shoot biomass in soils with a history of extreme drought (Figure 2). Range-expanding plants in general had the least shoot biomass compared to that of non-resident natives and alien plant invaders (Figure 2). The root biomass of invading plants was significantly greater in soils with a history of high native plant diversity contradicting the diversity-invasibility hypothesis (Table 1, Figure 3). Species-specific shoot and root biomasses of all invading plants used during the feedback phase are provided in supplementary figures 1 and 2.

**Table 1:** Effects of soil legacy of native plant diversity and extreme drought on the performance (shoot and root biomass) of invading plants of three categories (non-resident natives, range expanders and alien plants). The bold values indicate statistical significance (p-value<0.05). The degree of freedoms is estimated using Kenward-Roger approximations for mixed-effect models. Variance components are used as random intercept in mixed-effects models. Marginal R^2^ represents model fit with only fixed effects, whereas conditional R^2^ represents model fit with both fixed and random effects. sd stands for standard deviation. The upward arrows indicate increase of a given response variable (shown only for significant variables and those with

| Treatments (fixed effects) | Shoot biomass |  |  |  | Root biomass (log-transformed) |  |  |  |
| --- | --- | --- | --- | --- | --- | --- | --- | --- |
|  | F-value<sub>df</sub> | p-value | Marginal R<sup>2</sup> | Conditional R<sup>2</sup> | F-value<sub>df</sub> | p-value | Marginal R<sup>2</sup> | Conditional R<sup>2</sup> |
| Plant species richness (S) | 2.480<sub>1, 523.29</sub> | 0.115 |  |  | **4.747**<sub>1, 523.88</sub> | **0.029** ↑ |  |  |
| Extreme drought (D) | **7.104**<sub>1, 523.31</sub> | **0.007** ↑ |  |  | 0.101<sub>1, 523.77</sub> | 0.749 |  |  |
| Plant invader type (I) | **4.487**<sub>2, 14.51</sub> | **0.030** |  |  | 2.066<sub>2, 18.89</sub> | 0.154 |  |  |
| S×D | 0.081<sub>1, 523.38</sub> | 0.775 | 0.22 | 0.45 | 1.252<sub>1, 523.89</sub> | 0.263 | 0.06 | 0.24 |
| S×I | 1.742<sub>2, 523.38</sub> | 0.176 |  |  | 1.107<sub>2, 524.03</sub> | 0.331 |  |  |
| D×I | 0.647<sub>2, 523.32</sub> | 0.523 |  |  | 0.543<sub>2, 523.73</sub> | 0.581 |  |  |
| S×D×I | 1.245<sub>2, 523.36</sub> | 0.288 |  |  | 0.434<sub>2, 523.81</sub> | 0.647 |  |  |
| **Variance components** | **Variance (sd)** |  |  |  | **Variance (sd)** |  |  |  |
| Soil collection site | <0.001 (0.019) |  |  |  | <0.001 (0.027) |  |  |  |
| Plant species | 0.004 (0.069) |  |  |  | 0.080 (0.283) |  |  |  |

**Figure 2:**
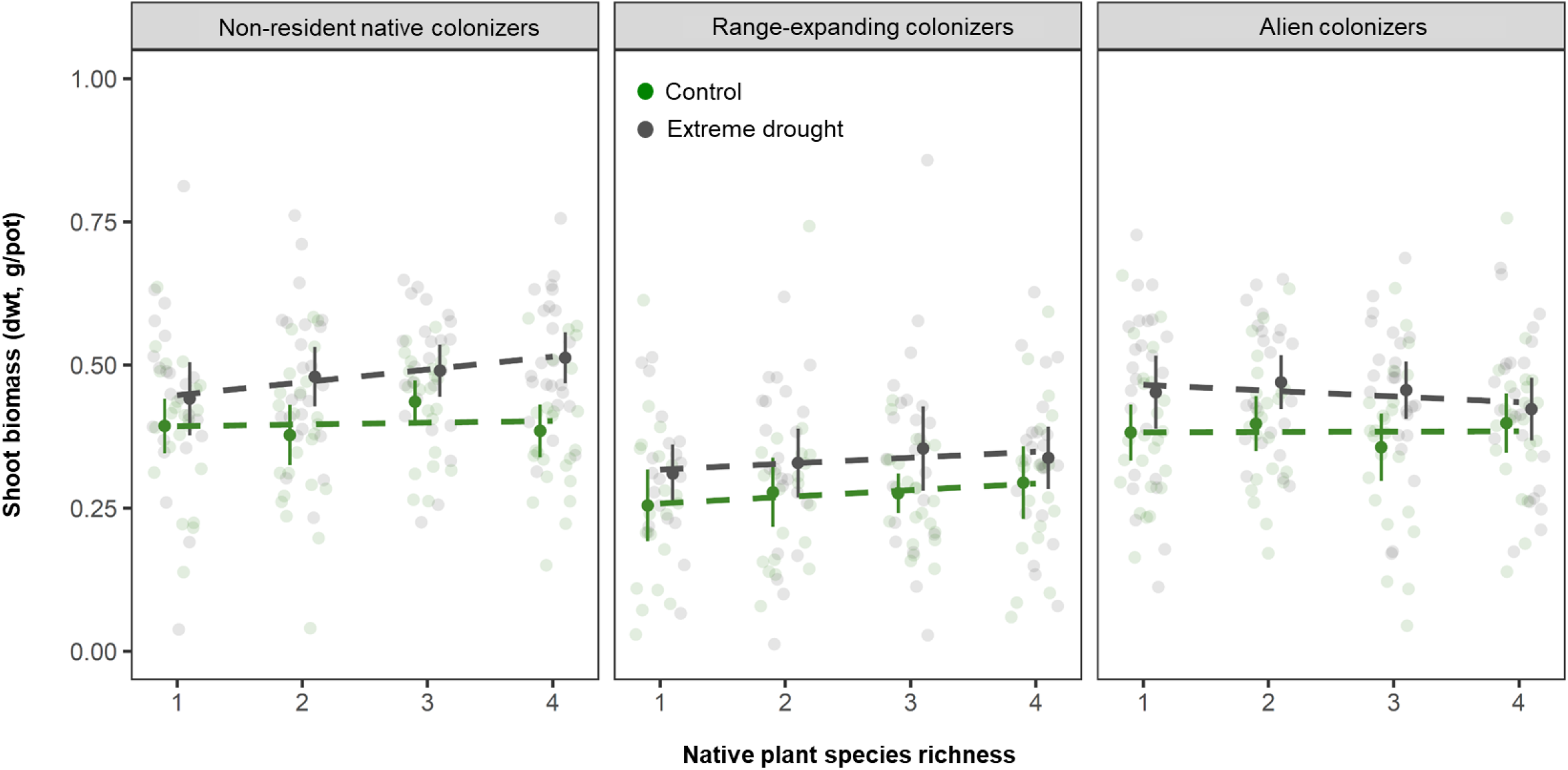
Feedback phase shoot biomass of three invader type plants (each group contained 4 plant species) in soils conditioned by native plant species richness and extreme drought. Control soils were with regular addition of water. Means are shown as dark colored points together with standard errors. Dashed lines indicated regression lines, whereas raw data points are shown as light colored points.

**Figure 3:**
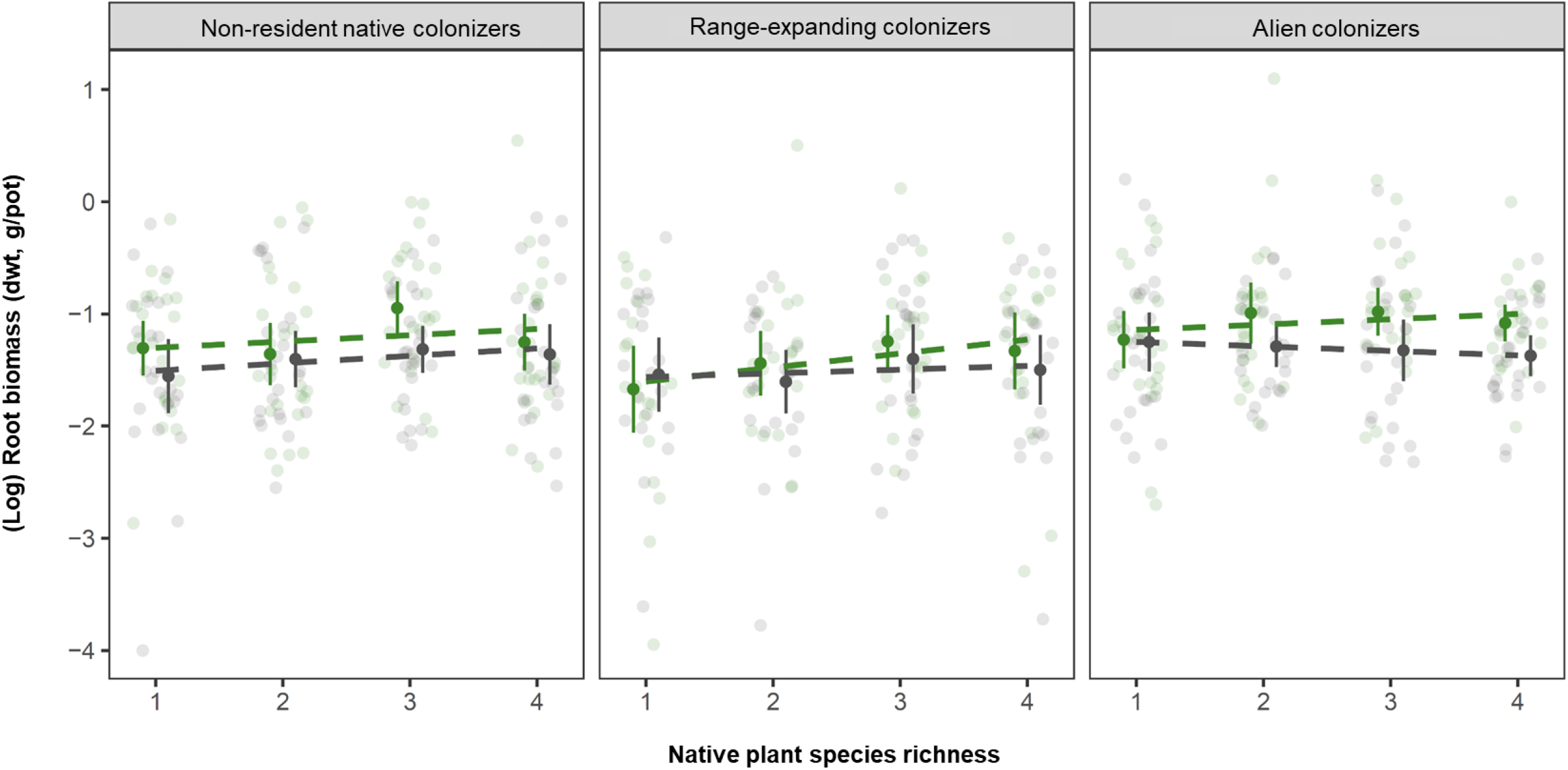
Feedback phase root biomass (log-transformed to meet linear model assumptions) of three groups of invading plants (each group contained 4 plant species) in soils conditioned by native plant species richness and extreme drought. Control soils were with regular addition of water. Means are shown as dark colored points together with standard errors. Dashed lines indicated regression lines, whereas raw data points are shown as light colored points.

To explore the underlying mechanisms that could have driven the effects of soil legacies of extreme drought and native plant diversity on the performance of invading plants during the feedback phase, we examined how abiotic (soil C, N and P responses) and biotic (native plant root and soil fungal responses) variables differed in soils from the conditioning phase. Extreme drought and native plant diversity consistently reduced native shoot and root biomass, whereas soil PO4 was also lower in soils with higher native plant diversity during the conditioning phase (Table 2). We quantified the response of 12 functional groups of soil fungi, of which the majority was influenced by native plant species richness (Table 2). However, only the relative abundance of total endophytic fungi (mainly the relative abundance of *Rhizoctonia solani*) decreased in soils that experienced extreme drought (Table 2). Multivariate analyses further showed significant compositional differences among soil fungal communities across native plant diversity (F-value1,175 = 29.36, p-value=0.01), whereas neither extreme drought nor its interaction with plant species richness significantly altered fungal community composition (p-value>0.05, Supplementary table 1).

**Table 2:**
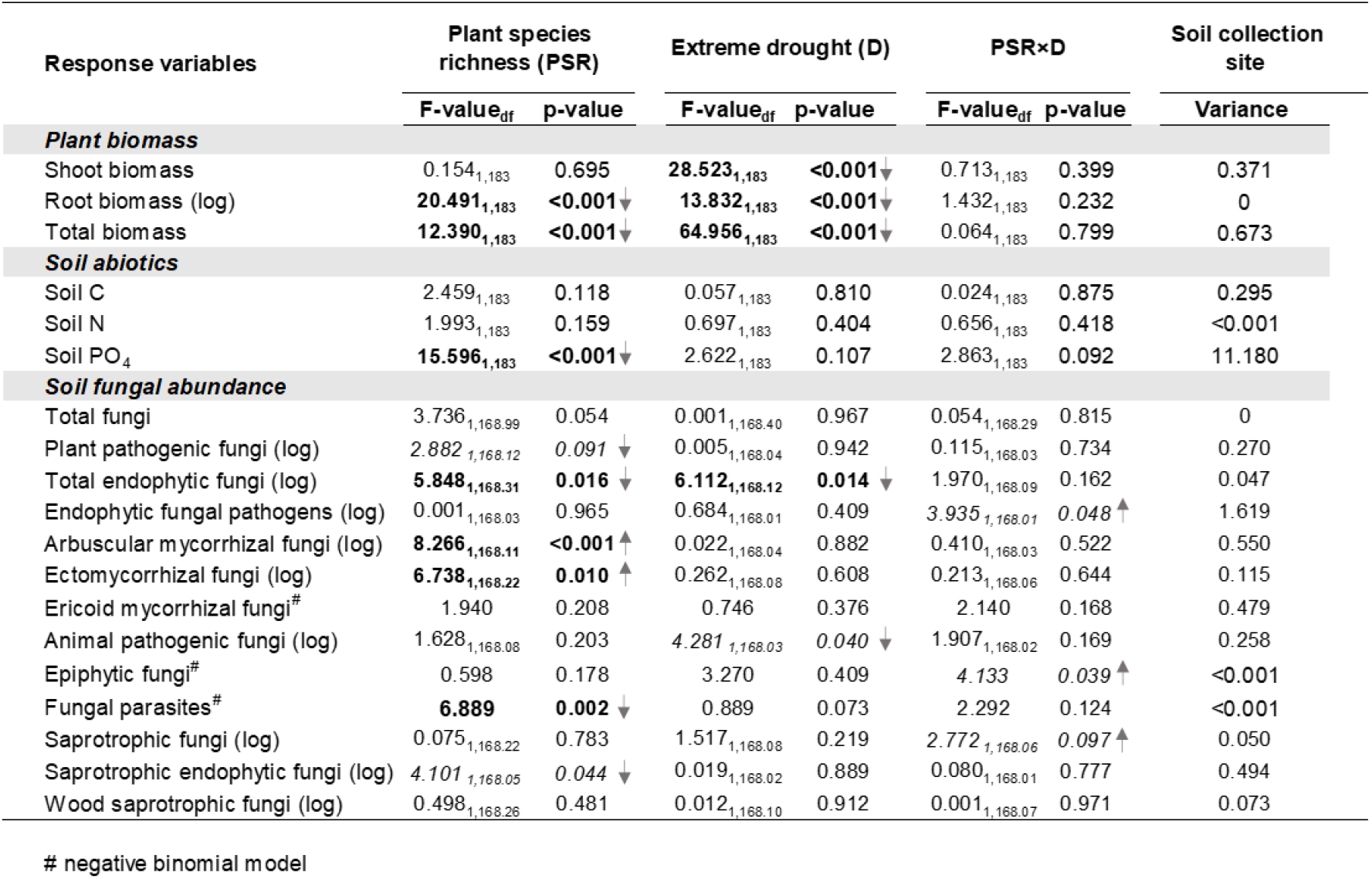
Effects of plant species richness, extreme drought and their interactions on plant biomass, soil abiotics and the abundance of soil fungal groups during the conditioning phase. The bold values indicate statistical significance (p-value<0.05) and italics are marginally significant (p-value<0.1). The upward and downward arrows indicate increase and decrease of a given response variable, respectively (shown only for statistically significant variables). Soil collection site was used as a random intercept in mixed models for which variance is reported. All mixed models were based on Gaussian error terms unless indicated by # for a given response variable (all other models included negative binomial error terms to account for overdispersion in the data).

| Response variables | Plant species richness (PSR) |  | Extreme drought (D) |  | PSR×D |  | Soil collection site |
| --- | --- | --- | --- | --- | --- | --- | --- |
|  | F-value<sub>df</sub> | p-value | F-value<sub>df</sub> | p-value | F-value<sub>df</sub> | p-value | Variance |
| ***Plant biomass*** |  |  |  |  |  |  |  |
| Shoot biomass | 0.154<sub>1,183</sub> | 0.695 | **28.523**<sub>1,183</sub> | **<0.001**↓ | 0.713<sub>1,183</sub> | 0.399 | 0.371 |
| Root biomass (log) | **20.491**<sub>1,183</sub> | **<0.001**↓ | **13.832**<sub>1,183</sub> | **<0.001**↓ | 1.432<sub>1,183</sub> | 0.232 | 0 |
| Total biomass | **12.390**<sub>1,183</sub> | **<0.001**↓ | **64.956**<sub>1,183</sub> | **<0.001**↓ | 0.064<sub>1,183</sub> | 0.799 | 0.673 |
| ***Soil abiotics*** |  |  |  |  |  |  |  |
| Soil C | 2.459<sub>1,183</sub> | 0.118 | 0.057<sub>1,183</sub> | 0.810 | 0.024<sub>1,183</sub> | 0.875 | 0.295 |
| Soil N | 1.993<sub>1,183</sub> | 0.159 | 0.697<sub>1,183</sub> | **0.404** | 0.656<sub>1,183</sub> | 0.418 | <0.001 |
| Soil PO<sub>4</sub> | **15.596**<sub>1,183</sub> | **<0.001**↓ | 2.622<sub>1,183</sub> | 0.107 | 2.863<sub>1,183</sub> | 0.092 | 11.180 |
| ***Soil fungal abundance*** |  |  |  |  |  |  |  |
| Total fungi | 3.736<sub>1,168.99</sub> | 0.054 | 0.001<sub>1,168.40</sub> | 0.967 | 0.054<sub>1,168.29</sub> | 0.815 | 0 |
| Plant pathogenic fungi (log) | 2.882<sub>1,168.12</sub> | 0.091 ↓ | 0.005<sub>1,168.04</sub> | 0.942 | 0.115<sub>1,168.03</sub> | 0.734 | 0.270 |
| Total endophytic fungi (log) | **5.848**<sub>1,168.31</sub> | **0.016** ↓ | **6.112**<sub>1,168.12</sub> | **0.014** ↓ | 1.970<sub>1,168.09</sub> | 0.162 | 0.047 |
| Endophytic fungal pathogens (log) | 0.001<sub>1,168.03</sub> | 0.965 | 0.684<sub>1,168.01</sub> | 0.409 | 3.935<sub>1,168.01</sub> | **0.048** ↑ | 1.619 |
| Arbuscular mycorrhizal fungi (log) | **8.266**<sub>1,168.11</sub> | **<0.001** ↑ | 0.022<sub>1,168.04</sub> | 0.882 | 0.410<sub>1,168.03</sub> | 0.522 | 0.550 |
| Ectomycorrhizal fungi (log) | **6.738**<sub>1,168.22</sub> | **0.010** ↑ | 0.262<sub>1,168.08</sub> | 0.608 | 0.213<sub>1,168.06</sub> | 0.644 | 0.115 |
| Ericoid mycorrhizal fungi<sup>#</sup> | 1.940 | 0.208 | 0.746 | 0.376 | 2.140 | 0.168 | 0.479 |
| Animal pathogenic fungi (log) | 1.628<sub>1,168.08</sub> | 0.203 | 4.281<sub>1,168.03</sub> | **0.040** ↓ | 1.907<sub>1,168.02</sub> | 0.169 | 0.258 |
| Epiphytic fungi<sup>#</sup> | 0.598 | 0.178 | 3.270 | 0.049 | 4.133 | **0.039** ↑ | <0.001 |
| Fungal parasites<sup>#</sup> | **6.889** | **0.002** ↓ | 0.889 | 0.073 | 2.292 | 0.124 | <0.001 |
| Saprotrophic fungi (log) | 0.075<sub>1,168.22</sub> | 0.783 | 1.517<sub>1,168.08</sub> | 0.219 | 2.772<sub>1,168.06</sub> | **0.097** ↑ | 0.050 |
| Saprotrophic endophytic fungi (log) | 4.101<sub>1,168.05</sub> | **0.044** ↓ | 0.019<sub>1,168.02</sub> | 0.889 | 0.080<sub>1,168.01</sub> | 0.777 | 0.494 |
| Wood saprotrophic fungi (log) | 0.498<sub>1,168.26</sub> | 0.481 | 0.012<sub>1,168.10</sub> | 0.912 | 0.001<sub>1,168.07</sub> | 0.971 | 0.073 |

We used path models to link the biotic and abiotic variables from the conditioning phase soils to explain the performance of invading plants during the feedback phase. We ran path models separately for the three types of invading plants, and only present those of native and alien plants (Figure 4) due to the lack of goodness of fit of the path model of range-expanding plants (details in method section). In none of the path models, we found soil N and soil P (soil PO4) from the conditioning phase explaining the performance of invader plants during the feedback phase, whereas root biomass production during the conditioning phase and the abundance of specific fungal groups did explain some of the variation in non-resident natives and alien plant performance. More specifically, the positive effects of extreme drought on invading non-resident native plants were mediated *via* lower root biomass production of resident native plants, which subsequently reduced the relative abundance of endophytic fungal pathogens (Figure 4 A). Given the negative relationship between endophytic fungal pathogens and invader performance, our path model results suggest that non-resident native plants are likely to experience weaker pathogenic pressure in soils that experienced extreme drought. Alien invader performance could also be partly explained through root biomass and fungal parasite (or mycoparasites: fungi that parasitize other fungi) responses during the conditioning phase (Figure 4 B). For instance, the positive soil legacy effect of native plant diversity on the performance of alien invaders (i.e., greater root biomass) was partly driven through the root biomass of native plants during the conditioning phase (Figure 4 B), whereas the performance of alien plants aboveground (i.e., shoot biomass) during the feedback phase was mediated through negative effects of native plant diversity on soil fungal parasites (Figure 4 B). Positive effects of extreme drought on alien invaders also occurred through decreased root biomass of resident native plants during the conditioning phase (Figure 4 B).

**Figure 4:**
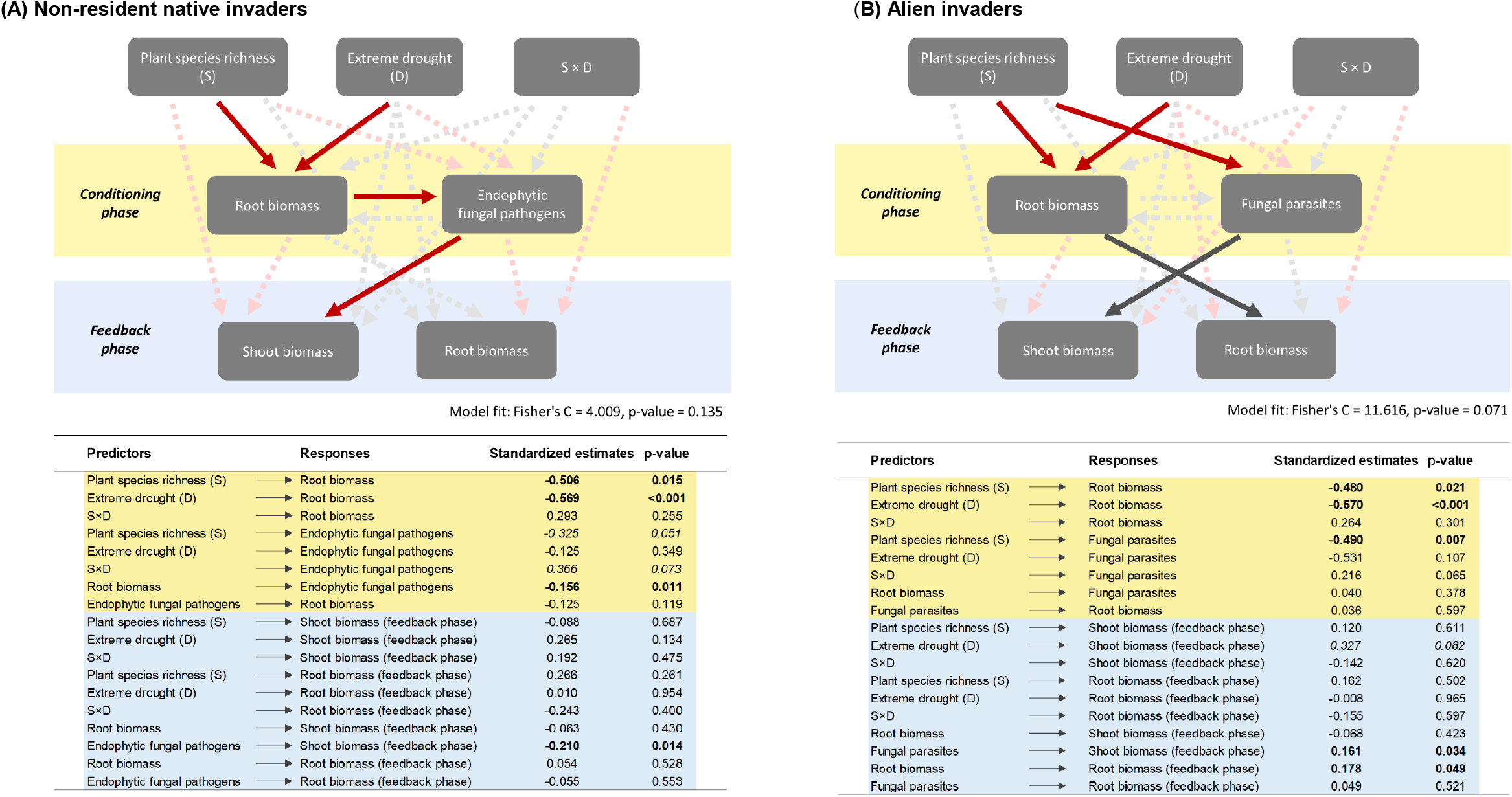
Results from path model analyses showing how soils conditioned by native plant diversity and extreme drought independently (lack of any interaction effect) affect the performance of (A) non-resident native invaders and (B) alien invasive plants *via* shifting root biomass (conditioning phase) and the abundance of soil fungal groups (conditioning phase). In the figure, thick lines (red: negative, gray: positive) represent statistically significant (p-values<0.05) relationships, whereas faded dashed lines represent statistically non-significant paths (p-value>0.05). The effect size (standardized path coefficients) and its statistical significance are tabulated below the figures. Bold numbers in the table are statistically significant, whereas italics numbers represent marginal significant effects (p-values less than 0.1 but greater than 0.05). Details of model structure are provided in the Material & Methods section, whereas model fit details are provided in the Result section. Path model for range-expanding plants with plant and fungal responses failed to have a good fit, and thus not shown here (Supplementary figure 4).

## Discussion

There is a growing consensus that the increasing intensity of drought events across the biosphere will alter plant community composition with detrimental consequences for native plant biodiversity and ecosystem functions^49,50^. Extreme droughts can shift plant community composition by benefitting one set of species while harming the others^51,52^, and thereby often enhancing plant turnover rates^53,54^. Many studies have suggested that alien plants have an advantage over native plants during and after extreme droughts^55–57^, but factors that could lead to such an advantage remain poorly understood. Through a plant-soil feedback experiment, we highlight that invading plants perform better in soils with an extreme drought legacy. This seemed true for all three types (non-resident natives, range-expanders, and alien plants) of invading plants irrespective of the soil legacy of native plant diversity. In fact, there were no signs of biotic resistance in soils with the legacy of native diverse community, but instead of a biotic facilitation through increments in root biomass production of invading plants. We identified some specific biotic pathways through which invading plants exploit soils of extreme drought or native plant diversity legacies. For instance, when extreme drought reduced the root biomass of native plant communities, it benefitted the non-resident native plants through a reduction in endophytic fungal pathogens. It is likely that lower production of roots during extreme droughts can simultaneously reduce the amount of fungal pathogens that reside inside the roots^58,59^.

Soil fungal communities are known to determine the success (or failure) of invading plants^60–62^. Endophytic fungi (those penetrating plant roots) in particular have received recent attention for their positive roles (e.g., enhanced nutrient uptake and pathogen defense) in driving the host performance including those of alien plants^60,63–65^, but also most pathogenic fungi that can inhibit plant performance are endophytic^66,67^. While we found that extreme drought and plant diversity both reduced the relative abundance of endophytic fungi during the conditioning phase (Table 2), it was the reduction of relative abundances of plant pathogenic endophytic fungi due to drought-induced lower root biomass during the conditioning phase through which extreme drought indirectly promoted the performance of non-resident native plants (Figure 4 A). Based on ITS sequencing data, it was pathogens like *Rhizoctonia solani* dominating the group of pathogenic endophytic fungi, which have a wide range of host plants^68^. It is well known that the high abundance of *R. solani* can severely damage their host plants^69,70^. A strong reduction of root biomass in our study during extreme drought could also have reduced the abundance of such endophytic fungi^71,72^, which subsequently promoted the performance of invading plants^73^. A recent study further suggested that fungal community responses to drought were closely related to root trait responses to soil water scarcity although pathogenic fungal responses were rather weak to drought^74^. We still know little about when and how (plant) host-associated pathogenic fungal species would respond to extreme droughts. The present results help us speculate that if some pathogens decline at the expense of lower resource availability (e.g., root biomass), invading plants can exploit such environments to establish themselves. Whether this mechanism further leads to a fast turnover of plant communities after extreme droughts remains to be tested.

Despite the overall weak soil legacy effects of native plant diversity relative to extreme droughts on the performance of all three types of invaders, our results still highlight biotic facilitation in promoting plant performance in these soils, specifically through an increase in root biomass of the invading plants. For the success of alien plants in native plant diversity soils, diversity-dependent reduction in mycoparasites (composed of basidiomycete taxa like *Chalciporus, Cystobasidium* and ascomycete taxa *Purpureocillium*) played some role (e.g., Figure 4 B). Mycoparasites are usually considered to regulate fungal pathogens and thereby promote the growth of plants^75^, but their decline could also be associated with increases of some other beneficial fungi that could foster the establishment of invading alien plants.

Both positive soil legacy effects of extreme drought and native plant diversity were in part driven by the reduction of root biomass production in resident native plants during the conditioning phase. Many previous studies have pointed to the importance of resource availability in the soil driving the success of invading plants^3,76^. Our results indeed suggest that enhanced performance of invading plants to partly associated with both resource availabilities in terms of root biomass and with the reduction of mycoparasites. The remaining roots of resident native plants in soils could affect nutrient availability through root decomposition, and further influence soil microorganisms that subsequently could determine the success of a new invading plant. We accordingly suggest that if an extreme drought lowers the root production in native plant communities and thereby alters soil fungal communities, it could open up opportunities for invading plants to establish, at least for non-resident native and alien plants used in our study. Indeed, our results are entirely based on soil legacy effects, and it is likely that the presence of native resident plants (for instance those that survive the extreme drought) could constrain the establishment of invading plants after extreme droughts through competitive interactions. Future studies can shed insight on what other aspects of plant roots (e.g., morphological and/or chemical traits) of native plant communities during extreme droughts and subsequent effects on soil fungal communities can influence the performance of invading plants.

A central question of 21^st^-century ecology is how multiple global change factors determine the dynamics of biodiversity change in ecosystems^77,78^. Extreme drought events are going to be a key driver of biodiversity dynamics, particularly through altering the plant- and soil community shifts in terrestrial ecosystems^79–82^. Our results confirm that the performance of invading plants enhances in soils with the legacy of extreme drought, and these effects may even exist independent of the background diversity of resident native plants. Moreover, we highlight that such drought legacies in soil are mediated through how extreme drought would alter the performance of previously present native plant communities and their associations with soil fungi.

## Materials and Methods

### Conditioning phase

The experiment consisted of two stages, the conditioning phase, and the feedback phase (Figure 1). The conditioning phase was performed to create the soil legacy of a native plant diversity gradient crossed with ambient water and extreme drought conditions. We grew four different native species to simulate different native plant diversity levels from 1 to 4 (Figure 1). These plant species were used based on their native status in Europe and their common co-occurrence in grasslands. Plant seeds were obtained from a local nursery company that collects seeds from the field (https://www.cruydthoeck.nl/). We added plants as seedling (about 2-3 cm in height) to gamma-sterilized soils (also collected from the field before the sterilization: Vlietberg, the Netherlands, 51° 86′ N, 5° 89’ E), which were then grown as monocultures, two species combination, three species combination, and four species combination - each containing four individuals as per the substitutive design. Live soils for the conditioning phase were collected where these species naturally grow from six different sites near Nijmegen, the Netherlands (close to the site where soils were collected for sterilization). These soils varied in their C, N and P content (Supplementary table 2), which is why we used soils from specific sites (even when they were in close vicinity) as random intercepts in mixed models (details below). We planted plant individuals in two-liter PVC pots (∼1800 g of sterilized soils mixed with 200 g of alive soil from the field, pot diameter 12.3 cm; pot height 13.0 cm) during the conditioning phase with all diversity levels replicated 24 times to create an adequate amount of soils for the feedback phase. Moreover, the field soil inoculation (from six sites) to these pots was replicated four times which led to 96 pots for native diversity treatments. We then doubled the diversity gradient pots (96) to create both ambient and extreme drought treatments. Please note that given the substitutive design, we were not able to replicate all compositions of plant pairs and three species combinations, and our statistical analysis is based on the diversity levels (1, 2, 3 and 4), which were adequately replicated during the conditioning phase.

Moisture levels during the conditioning phase were monitored regularly by weighing the pots and correcting when large differences (e.g., exceeding a difference of ∼10 g) within the treatments occurred. The conditioning of the soil with a native plant diversity gradient took place for ten weeks in total (eight weeks with a regular supply of water, and the last two weeks with half of the pots receiving no water at all). This extreme drought treatment resulted in a complete wilting of plants at the end of the conditioning phase (Figure 1). At the end of the conditioning phase, the moisture percentage in the soil was calculated using the gravimetric method to evaluate soil water differences between ambient and extreme drought conditioning treatments. The extreme drought treatment resulted in four times less soil moisture compared to the ambient conditions, which was sufficient to create wilting in plants across diversity treatments (Figure 1 b, Supplementary figure 3).

At the end of the conditioning phase (week 10 of the plant growth), plant shoots were clipped, roots were collected from half of the conditioned soil, and some of the conditioned soils were kept for the measurement of soil abiotic and fungal analysis. The roots were washed thoroughly to ensure no influence of residual soil on the root biomass. Both roots and shoots were dried at 40 °C for several days and weighed afterwards for the estimation of the shoot and root dry biomass. Soil samples (approximately ten grams) were taken from the conditioned to measure soil C, N and P content. Amounts of carbon (C) and nitrogen (N) were determined using an elemental analyzer (Flash EA 1112, Thermo Scientific) by following the Micro-Dumas combustion method. We further determined the amount of phosphorous (P) by extracting NaHCO3 and determining the plant-available orthophosphates in the soil following the Olsen P method ^83^.

### Fungal groups measurements

We measured the relative abundance of various soil fungal groups from the conditioning phase soils by amplicon sequencing using the Illumina Miseq PE300 platform at BGI (Hong Kong). Total DNA was extracted from 250 mg of fresh soil using the DNeasy Powersoil kit (Qiagen, Hilden, Germany) according to manufactures protocol. For multiplexing and re-identification individual samples adapters and barcodes were added to samples by polymerase chain reaction (PCR). We used the primers ITS4 and ITS9 targeting the ITS2 region^84^. PCRs were performed in 25-μl reaction mixtures and contained 2.5 μl dNTP (4005 μM), 0.15 μl of FastStart Expand High Fidelity polymerase (Roche Applied Sciences), 2.5 μl 10× PCR buffer with MgCl2, 1μl MgCl2 (25mM), 1.25 μl BSA (4mg/ml), 0.5 μl of each of the two primers (10mM) and 1 μl soil DNA (∼5–20 ng). The PCR conditions involved initial step at 95°C for 5 min, followed by 35 cycles of 95°C for 45 seconds and 54°C for 60 seconds, and then 72°C for 90 seconds. The final extension step was 72°C for 10 minutes. The final PCR product was purified using Agencourt AMPure XP magnetic bead system (Beckman Coulter Life Sciences, Indianapolis, Indiana, USA) with a volume ratio of PCR product to beads of 1: 0.7. After purification, we analyzed the PCR products in a Fragment Analyzer using a Standard Sensitivity NGS Fragment Analysis kit (1bp-6000bp) and following manufacturer’s instructions (Advanced Analytical Technologies GmbH, Heidelberg, Germany). The purified amplicons were diluted to the same concentration before sending for sequencing using the Illumina MiSeq platform for 300 bp paired-end reads.

ITS sequences were analyzed using the PIPITS pipeline (version 2.7, standard settings, Gweon et al. 2015) with VSEARCH implemented to pair sequences. The ITS2 region was extracted using ITSx ^86^ and short reads (<100bp) were removed. Chimeric sequences were removed by comparing with UNITE uchime database (version 8.2). Clustering of reads into OTUs was performed at an identity threshold of 97%. Sequences were aligned and classified using with RDP against the UNITE fungal database^87,88^ to obtain a match with the closest known species for each OTU. The resulting OTUs were parsed against the FunGuild (v1.1) database to assign putative life strategies^89^ for the OTUs with ‘probable’ or ‘highly probable’ classification and the assignment was further curated using in-house databases^90^. We resolved 12 major fungal functional groups that were used in further analysis. These groups were 1) plant pathogenic fungi, 2) total endophytic fungi, 3) endophytic fungal pathogens, 4) arbuscular mycorrhizal fungi, 5) ectomycorrhizal fungi, 6) ericoid mycorrhizal fungi, 7) animal pathogenic fungi, 8) epiphytic fungi, 9) fungal parasites, 10) total saprotrophic fungi, 11) saprotrophic endophytic fungi, 12) wood saprotrophic fungi.

### Feedback phase

To prepare for the feedback phase, the conditioned soils (300 grams) were mixed with gamma-sterilized soils (300 grams)- these soils were the same that were collected from the field from Vlietberg, the Netherlands (51° 86′ N, 5° 89’ E) for the conditioning phase. One-liter PVC pots (width 10 cm; length 10 cm; height 11 cm) were planted with an individual plant (Figure 1). Seeds of feedback phase plants were also purchased from the same local nursery company (https://www.cruydthoeck.nl/) where we purchased seeds for the conditioning phase plants, except that *Ceratochloa sitchensis* (the alien Poaceae) and *Senecio inaequidens* (the alien Asteraceae) were collected from the field in the Netherlands (near Wageningen). List of all 12 feedback-phase plant species is provided in Figure 1. These commonly found grassland plants were chosen in the feedback phase to create a phylogenetically balanced design relative to the conditioning phase (Figure 1). Each of these twelve species was planted after germination (about 2-3 cm in height) to the feedback phase pots filled with conditioned soils with a history of different native plant diversity levels (one to four species) and drought treatments. We grew each of the 12 feedback species separately to examine their performance in the conditioned soils. Each feedback species was replicated six times per drought and diversity treatments, which were distributed evenly across six soil origins from the conditioning phase. In this way, we examined the performance 576 plant individuals during the feedback phase. The plants were grown for six weeks and were watered regularly. The moisture difference between the extreme drought soils and ambient soils was overcome by adding 36 ml extra water to extreme drought soils at the beginning of the feedback phase. In this way, we excluded the effects of moisture difference already at the start of the feedback phase. After six weeks, the shoots were harvested by clipping, dried (again at 40 °C for several days) and weighed. Roots were harvested by washing them out of the soil gently and carefully prior to drying (also at 40 °C for several days) and weighing.

### Statistical analysis

Both conditioning and feedback phase response variables were analyzed using mixed-effects models. For the feedback phase, we explained variation in invader plant performance (shoot and root biomass) using a three-way interaction model with native plant diversity, drought, and invader type as three fixed effects, whereas soil inoculum was used as random intercept with six different levels (i.e., six different soils). Additionally, we used specific plant species as another random intercept in our feedback phase mixed-effect models (Table 1). Given that we measured many variables during the conditioning phase to understand soil biotic and abiotic characteristics as well as plant performance, we ran models separately for all those variables (Table 2). For all these mixed-effect models, six soil inoculums were again used as a random intercept, whereas native plant diversity and drought were the fixed effects. The list of all the response variables during the conditioning phase is provided in table 2, and could be categorized into plant community, soil abiotic variable and soil fungal group responses. Linearity assumptions of the model were visually inspected using the homogeneity of variance, and when not met, we either log-transformed the data or used negative binomial error distribution instead of normal distribution (indicated in Table 2).

Finally, we examined the relationship between the soil legacy treatments and the performance of invading plants in the feedback phase using path models to be able to distinguish between direct and indirect effects from the conditioning phase treatments that influence the feedback phase invading plants. Model structures in our path models were similar to that of mixed-effect models for conditioning and feedback phase response variables. However, since we ran path models separately for three types of invading plants, we excluded specific plant species as a random term. Indeed, soil origin having six levels compared to four levels of plant species as a random intercept within the invader type provides a more robust parameter estimation in mixed models ^91^. Moreover, two independent random intercepts (soil type and plant species) in mixed-effect models prevented model convergence, which is why we opted to use a single random term instead of two in all our path models. We included soil fungal and plant response data from the conditioning phase and linked them to invading plant performance (shoot and root biomass) during the feedback phase. For some variables, we tested both pathways to identify the causal links (Figure 4). Model selection was used based on the inclusion of variables that resulted in non-saturated path models e.g., model overfit) as per Shipley’s test of d-separation which yields Fisher’s C statistic (Chi-square distributed)^92^. None of our path models showed a good fit (p-value <0.05) when ran for range-expanding invaders with similar set of variables (e.g., soil fungal groups and/or soil nutrients) used for local native and alien invasive plants. The inclusion of soil nutrients in path models also failed to provide a good fit (p-value<0.05) for non-resident native and alien plants. We eventually selected two path models-one for non-resident and one for alien invaders with plant biomass responses and one of the fungal group responses to associate with the respective performance (both shoot and root biomass) of invading plants (Figure 4). Moreover, the path model for the alien plant was further reduced to obtain a good fit (the direct paths from conditioning phase plant diversity to feedback phase plant shoot and root biomass were removed), although for the sake of comparison between local and alien invaders, we have shown the full model for alien plants as well (Figure 4).

All statistical analyses were carried out in R statistical software version 4.1.0 ^93^. Mixed effect models were run with the lme4 package for R, whereas we used the performance package to check the linearity model assumption^94^. The F-value, p-value and degrees of freedom were estimated using the lmerTest package^95^. Multi-variate analysis of fungal communities were performed using the *manyglm* function from the mvabund package which incorporates the framework of generalized linear models ^96^. In path models, we used the nlme package^97^ to run the mixed-effect models. The path model analysis was carried out with the piecewiseSEM package^98^.

## Acknowledgements

We thank Jake Alexander and Arjen Biere for their suggestions on the experimental design. We are grateful to Ludovico Formenti for his help in drawing figure 1. MPT acknowledges the funding from the Swiss State Secretariat for Education, Research and lnnovation (SERI) under contract number M822.00029. WHvdP acknowledges the support from ERC advanced grant (ERC-ADV 323020 SPECIALS).

## Author contribution

MPT and WHvdP conceived the study. MvdS, MPT, RAW, SEH, FtH, SG, CQ, KS performed the experiment. MPT analysed the data with inputs from SEH and wrote the manuscript together with WHvdP. All authors contributed to revisions.

## Competing Interest Statement

The authors declare no conflict of interest.

## Data availability

All data used for the analysis is uploaded as a supplementary data file.

## Notes

### Competing Interest Statement

The authors have declared no competing interest.

### Summary of Updates

Figure 1 and Figure 4 were not properly uploaded in the manuscript file. They have been correctly uploaded in this version.

