## Supplementary Table 1-2, Supplementary Figures 1-4 for "Soil legacies of extreme droughts enhance the performance of invading plants"

**Supplementary table 1:** Effects of plant species richness and extreme drought on soil fungal composition during the conditioning phase based on multivariate generalized linear models. Fungal data in this analysis were included as relative abundances of specific fungal groups. The bold values are statistically significant (p-value<0.05). The df stands for degree of freedom.

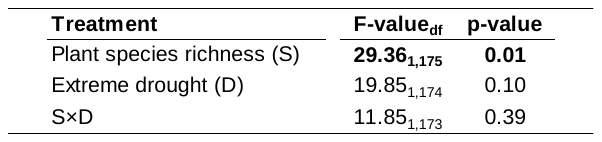

**Supplementary table 2:** C, N and PO4 content of six different soils that were used as an inoculum for soil conditioning phase. These soils were collected from the six different closed by areas where native plants used in our study co-occur.

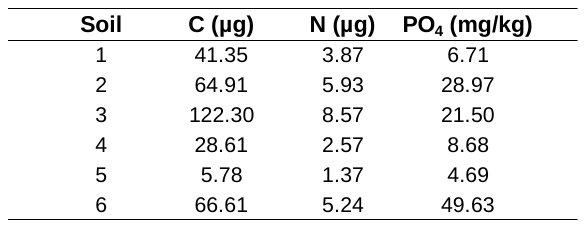

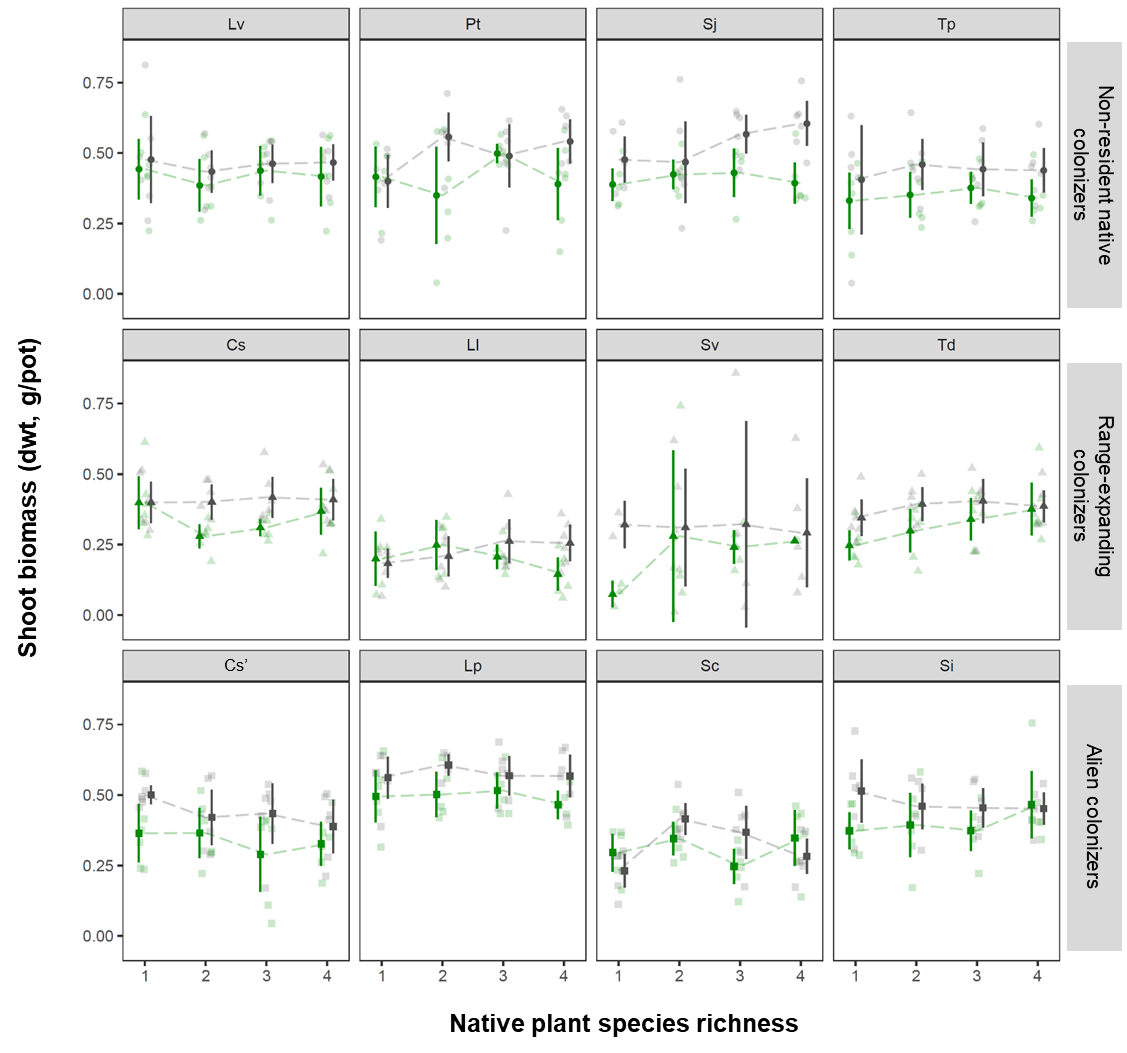

**Supplementary figure 1:** Shoot biomass of specific plants used during the feedback phase. Control water treatments are shown in green, whereas extreme drought treatments are shown in gray. Lv: *Leucanthemum vulgare*; Pt: *Poa trivialis*; Sj: *Senecio jacobaea*; Tp: *Trifolium pretense*; Cs: *Centaurea stoebe*; Ll: *Lathyrus latifolius*; Sv: *Setaria verticillata*; Td: *Tragopogon dubius*; Cs’: *Ceratochloa sitchensis*; Lp: *Lupinus polyphyllus*; Sc: *Soliddago canadensis*; Si: *Senecio inaquidens.*

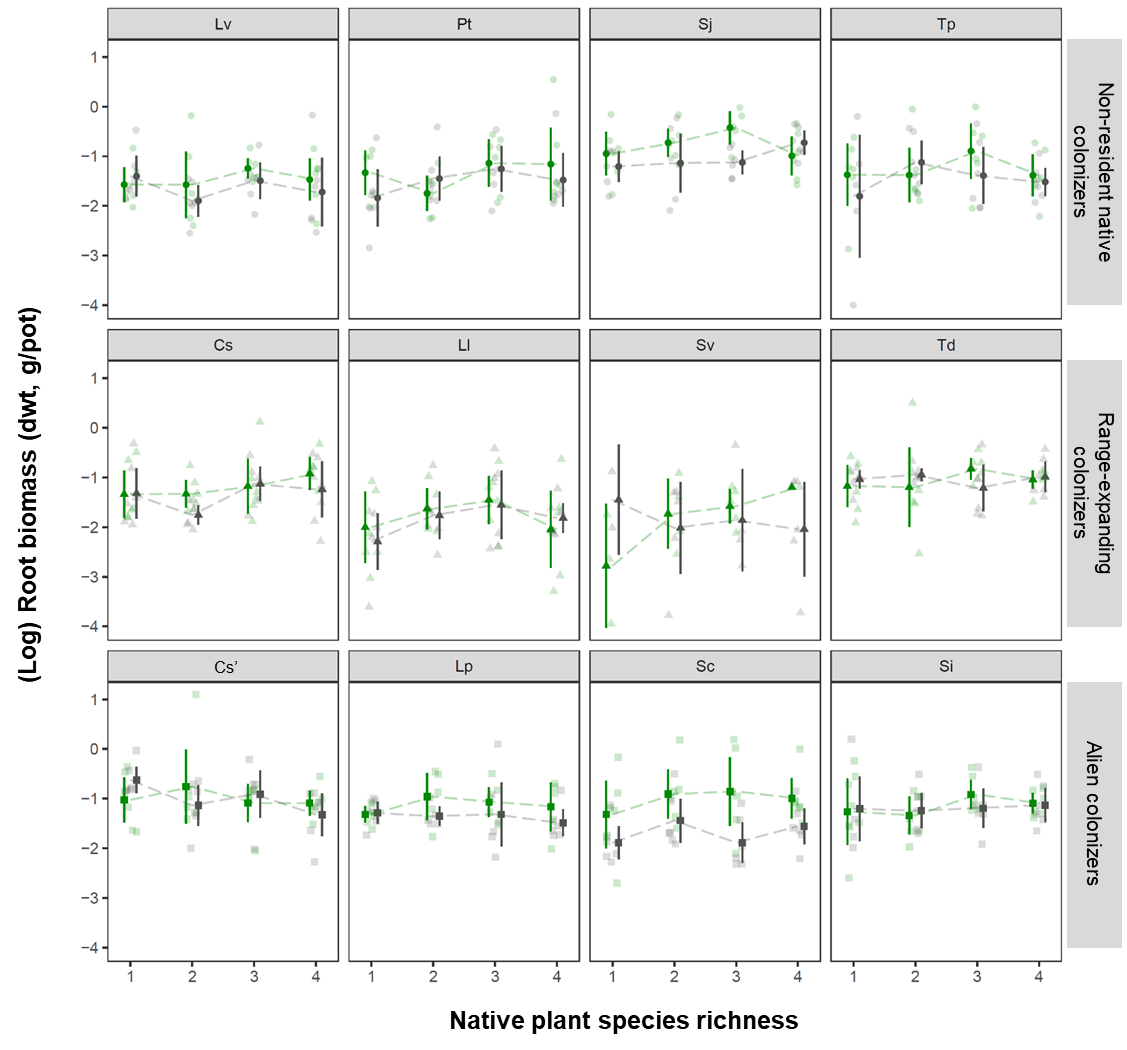

**Supplementary figure 2:** Root biomass of specific plants used during the feedback phase. Control water treatments are shown in green, whereas extreme drought treatments are shown in gray. Lv: *Leucanthemum vulgare*; Pt: *Poa trivialis*; Sj: *Senecio jacobaea*; Tp: *Trifolium pretense*; Cs: *Centaurea stoebe*; Ll: *Lathyrus latifolius*; Sv: *Setaria verticillata*; Td: *Tragopogon dubius*; Cs’: *Ceratochloa sitchensis*; Lp: *Lupinus polyphyllus*; Sc: *Soliddago canadensis*; Si: *Senecio inaquidens.*

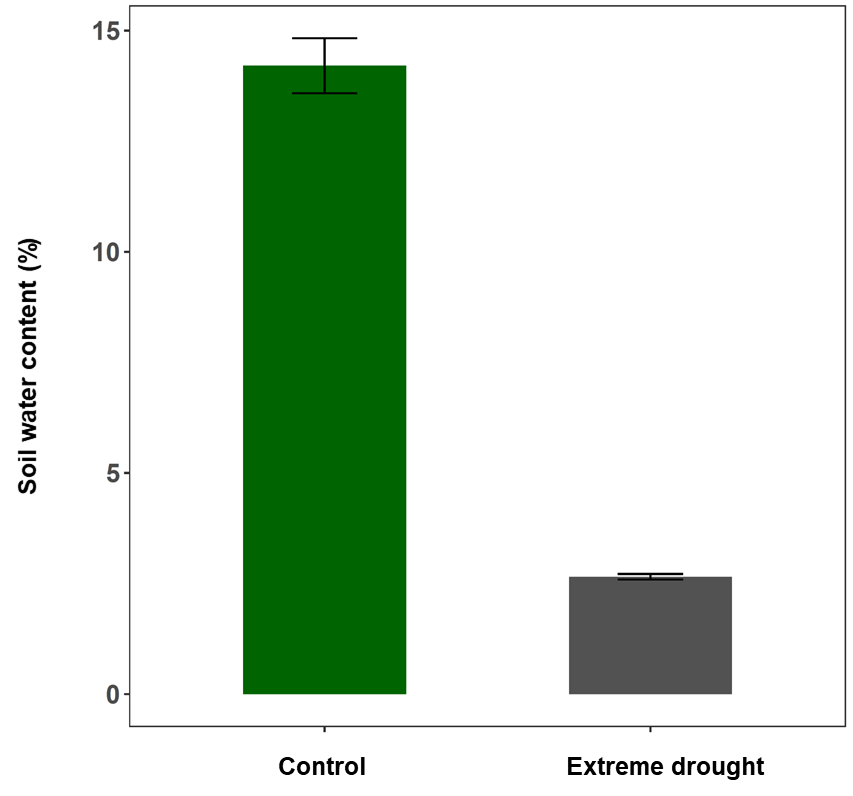

**Supplementary figure 3:** Difference in soil water content between our control experimental units and extreme drought units during the conditioning phase. Soil water content was determined by calculating the difference between the weight of the soil before air drying and the weight of the soil after air drying (at 70°C for a week).

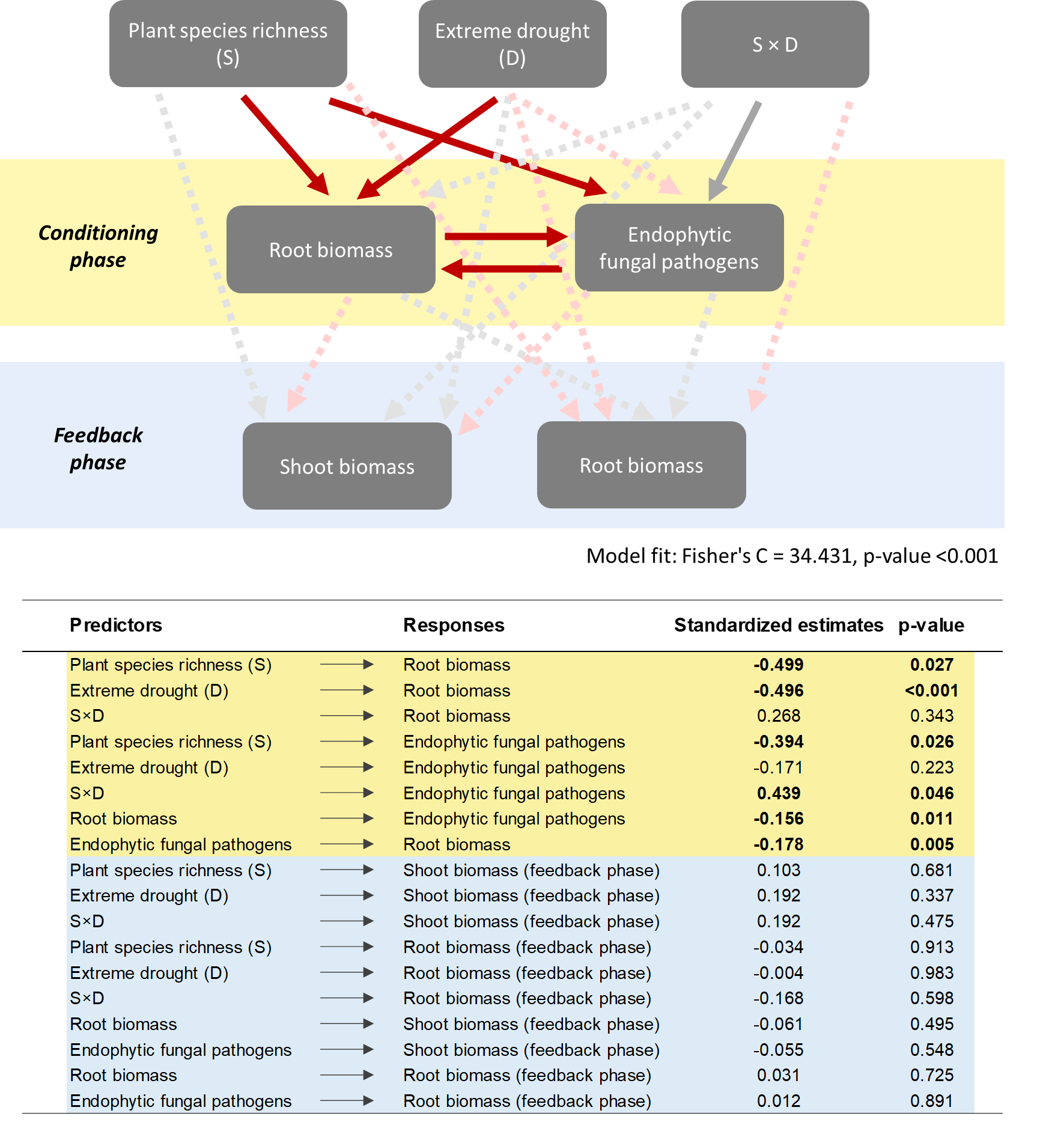

**Supplementary figure 4:** Results from path model analysis showing how soils conditioned by native plant diversity and extreme drought independently affect the performance of range-expanding plants. Given that the path model with range-expanding plants failed to meet the goodness of model fit criteria of Fisher C test (p-value > 0.05), these results were not discussed in the main text.
